# Eukaryotic transposable elements as “cargo carriers”: the forging of metal resistance in the fungus *Paecilomyces variotii*

**DOI:** 10.1101/2020.03.06.981548

**Authors:** Andrew S. Urquhart, Nicholas F. Chong, Yongqing Yang, Alexander Idnurm

**Affiliations:** School of BioSciences, the University of Melbourne, VIC, 3010, Australia; Inner Mongolia Academy of Agricultural & Animal Husbandry Sciences, Hohhot 010031, China

**Keywords:** Eurotiales, gene cluster, metal homeostasis, horizontal gene transfer, transposon

## Abstract

The horizontal transfer of large gene clusters by mobile elements is a key driver of prokaryotic adaptation in response to environmental stresses. Eukaryotic microbes face similar environmental stresses yet a parallel role for mobile elements has not yet been established. A stress faced by all microorganisms is the prevalence of toxic metals in their environment. In fungi, identified mechanisms for protection against metals generally rely on genes that are dispersed within an organism’s genome. Here we have discovered a large (∼85 kb) region that confers resistance to several metals in the genomes of some, but not all, strains of a fungus, *Paecilomyces variotii*. We name this region *HEPHAESTUS* (*Hϕ*) and present evidence that this region is mobile within the *P. variotii* genome with features highly characteristic of a transposable element. While large gene clusters including those for the synthesis of secondary metabolites have been widely reported in fungi, these are not mobile within fungal genomes. *HEPHAESTUS* contains the greatest complement of host-beneficial genes carried by a transposable element in eukaryotes. This suggests that eukaryotic transposable elements might play a role analogous to their bacterial counterparts in the horizontal transfer of large regions of host-beneficial DNA. Genes within *HEPHAESTUS* responsible for individual metal resistances include those encoding a P-type ATPase transporter, PcaA, required for cadmium and lead resistance, a transporter, ZrcA, providing resistance to zinc, and a multicopper oxidase, McoA, conferring resistance to copper. Additionally, a subregion of *Hϕ* conferring resistance to arsenate was identified. The presence of a strikingly similar cluster in the genome of another fungus, *Penicillium fuscoglaucum*, suggests that *HEPHAESTUS* arrived in *P. variotii* via horizontal gene transfer.

## INTRODUCTION

Metals are used within diverse fields such as agriculture, construction, electronics and pharmaceutical science. A number of these, such as cadmium, lead and arsenic, are toxic even at low levels of exposure and thus environmental contamination resulting from their use is an increasing concern ^1^. The prevalence of toxic metals in the environment is of particular relevance to fungal biology. Some fungi are able to accumulate metals to high concentrations, making such fungi dangerous for human consumption ^2,3^, allowing bioremediation of contaminated sites ^4^ or concentrating valuable metals for extraction ^5^. Zinc and copper play essential roles in many cellular functions and are often important in fungal virulence ^6^. However, at high concentrations these metals can also be toxic; for example, copper-based molecules are frequently used as fungicides in agriculture.

Compared to prokaryotes, our understanding of the evolutionary mechanisms through which eukaryotes have adapted to environmental stresses – such as to metal toxicity – is relatively limited. One fungal example is the recent expansion of the *CUP1* genes encoding metallothioneins in wine and sake-producing strains of *Saccharomyces cerevisiae*. This gene expansion is hypothesized to be a response to the use of copper-based materials during fermentation and/or copper sulfate fungicides in vineyards ^7,8^.

In prokaryotes, horizontal transfer of integrative and conjugative elements (ICEs) has played a key role in the evolution of heavy metal tolerance. ICEs are mobile pieces of DNA that are inserted into the host genome and encode the genes necessary for conjugation thus allowing their spread between bacterial hosts. These elements frequently carry additional host-beneficial cargo genes facilitating the transfer of key traits such as resistance to antibiotics. There are several examples of ICE elements carrying the genes required for tolerance to heavy metals; for example, tolerance to copper in *Pseudomonas syringae* ^9^, copper and arsenic in *Salmonella enterica* ^10^, and copper and zinc in *Histophilus somni* ^11^. The acquisition of ICEs is thus a means by which bacterial strains can evolve resistance to heavy metals. The potential role of horizontal gene transfer events in the evolution of metal tolerance has not been explored in eukaryotes. However, horizontal gene transfer, while less extensive than in prokaryotes, is now recognized as an important evolutionary force in other areas of eukaryote biology including fungal virulence and nutrient utilization ^12,13^.

The horizontal transfer of multigene traits is facilitated by the fact that filamentous fungi feature distinct regions in which genes associated with the same process are physically clustered, especially for the synthesis of secondary metabolites ^14-16^. Other gene clusters play roles in primary metabolism such as nitrate assimilation ^17^ and biotin synthesis ^18^. Small clusters of genes involved in metal homeostasis such as iron uptake and arsenic detoxification have been identified in fungi ^19^. In *S. cerevisiae* three linked genes mediate resistance to arsenicals ^20^. How such clusters are transferred between species is not understood. Although it has recently been hypothesized that gene clusters are horizontally transferred between eukaryotic species on large transposable elements ^21,22^, transposons carrying large regions of host-beneficial DNA have not been reported.

*Paecilomyces variotii* is a cosmopolitan member of the Eurotiales that is capable of growing under a wide range of environmental conditions. For example, it is noteworthy for its high level of resistance to heat which causes it to be a frequent cause of food spoilage in heat treated foods ^23,24^. It has also been isolated from diverse environments including as a plant endophyte ^25^, marine sources ^26^, human infections ^27^, and indoors. The ecology of this species is currently poorly understood.

Recently, a genetic transformation system has been developed and the genomes of three *P. variotii* strains have been sequenced ^28,29^. Here, we describe the discovery and functional analysis of part of the genome, absent from some strains, that features genes encoding proteins involved in metal detoxification or protection. This gene cluster is distinct from previously described clusters associated with heavy metal resistance because it encodes > 30 genes and provides resistance to at least five different metals: arsenic, cadmium, copper, lead and zinc. Furthermore, we present evidence that this region is mobile within the *P. variotii* genome and may have arisen through horizontal gene transfer, suggesting a role for transposable elements in the transfer of large genomic regions conferring beneficial traits between fungi.

## RESULTS

### The *HEPHAESTUS* region is only present in certain *Paecilomyces variotii* strains

Comparison of two *P. variotii* strains revealed a region of the genome present within strain CBS 144490 but absent from CBS 101075 (Figure 1a). Examination of the predicted genes within this region revealed that many have similarities to those involved in protection against various toxic metals (Supplementary Table 1). We name this genetic element *HEPHAESTUS* (*Hϕ*), after the Greek god of metallurgy. Genes within the cluster were annotated based on sequence similarity to homologs in other fungi. Sequencing four additional *P. variotii* strains revealed the presence of the cluster in strains FRR 1658, FRR 3823 and FRR 5287, and its absence from strain FRR 2889 (Figure 1b). A recently sequenced strain of *P. variotii*, MTDF-01, also lacks this region ^28^. Neither of the other published *Paecilomyces* genomes, *P. formosus* strain number 5 ^30^ or *Paecilomyces niveus* strain CO7 ^31^, have the cluster. “*Byssochlamys”* (anamorph *Paecilomyces*) strain BYSS01 ^32^ has part of the cluster, but this genome appears to represent a *Monascus* species rather than *Paecilomyces* based on BLAST searches of phylogenetic markers (data not shown).

**Fig. 1.**
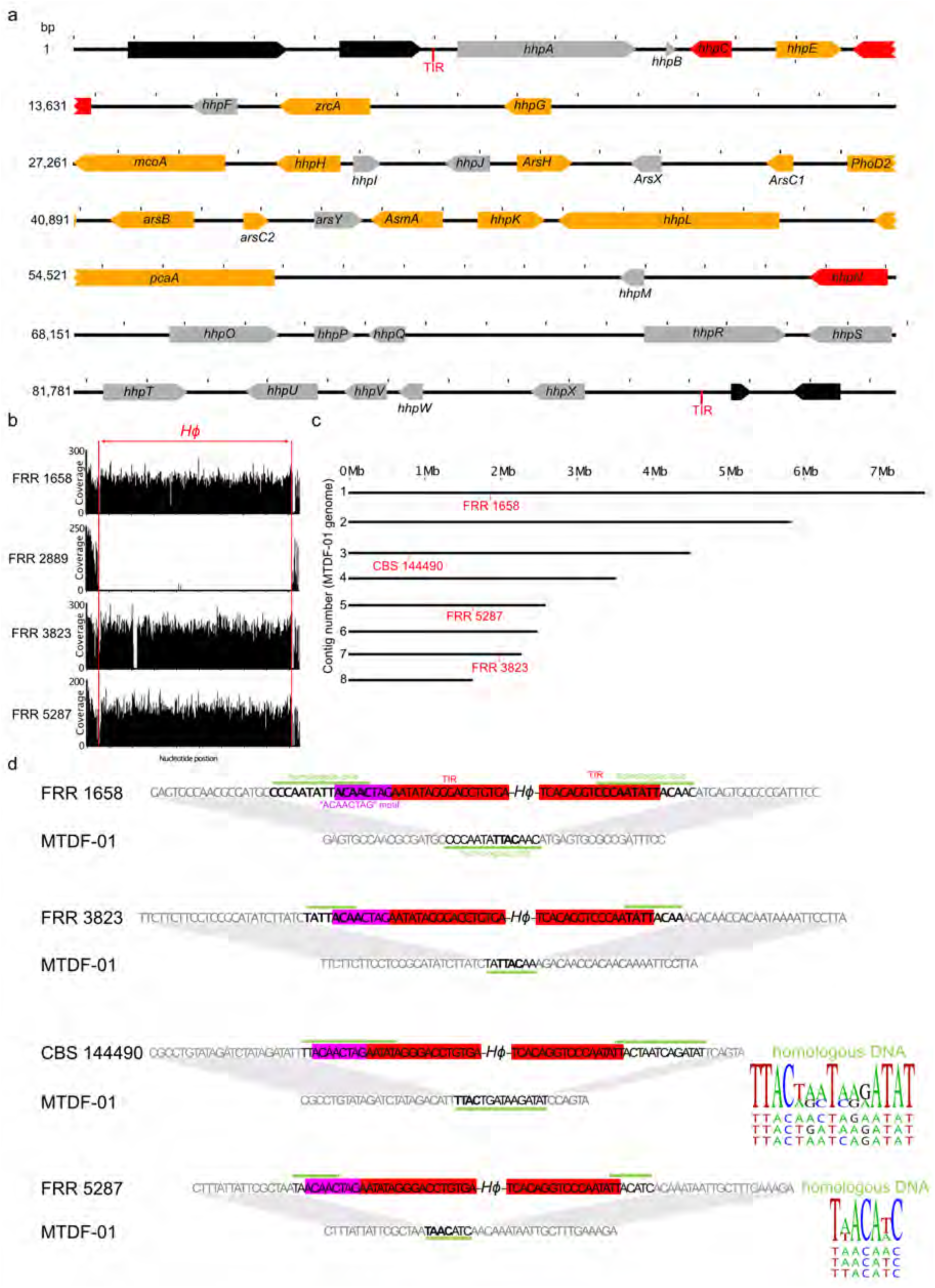
*HEPHAESTUS (Hϕ*), a large gene cluster in *P. variotii*. **a**, Map of the *HEPHAESTUS* region and flanking DNA in *P. variotii* CBS 144490. Genes with putative roles in metal resistance based on homology are highlighted in orange. Genes with similarity to those involved in transposition of transposons are highlighted in red. Flanking genes conserved in the *Hϕ-*strain CBS 101075 in black. TIR; terminal inverted repeats. Tick marks represent 1 kb of nucleotide sequence. **b**, Read mapping to the *Hϕ* region shows that of the four newly sequenced strains FRR1658, FRR 3823 and FRR 5287 contain *Hϕ* but FRR 2889 does not. **c**, Genomic location of *Hephaestus* in four strains annotated onto the 8 largest contigs of the *P. variotii* strain MTDF-01 assembly. Three other sequenced strains, CBS 101075, FRR 2889 and MTDF-01, do not contain *Hϕ*. **d**, Sequences of the insertion sites in the four *Hϕ*+ *P. variotii* strains. Features include the terminal inverted repeats (TIR; red), the “ACAACTAG” motif (purple), the TTAC/TAAC sequence (bold) and regions of microhomology (green).

### *HEPHAESTUS* is a mobile element within the *Paecilomyces* genome

The edges of the cluster feature an inverted repeat AATAT[A/T]GGGACCTGTGA, where the [A/T] is different between the two copies. Furthermore, one gene encodes a Tc-1 like transposase, another contains an RNase H domain homologous to that found in reverse transcriptases, and there are also remnants of other transposable elements. To explore the possibility that *Hϕ* is a mobile element we compared the genomic location of *Hϕ* within the four sequenced *P. variotii* strains that have the cluster (*Hϕ+* strains). Each contained *Hϕ* at an independent location on a different assembly contig (Figure 1c). Examination of the *Hϕ* integration sites revealed clear patterns: *Hϕ* appears to have been inserted into either a TTAC site (in FRR 1658, FRR 3823, CBS 144490) or a TAAC site (in FRR 5287) (Figure 1d). In each case the target site showed microhomology to the edges of *Hϕ* suggesting that microhomology is important for *Hϕ* transposition. A further distinctive feature was the presence of the “ACAACTAG” motif next to the right terminal repeat. We hypothesize that this sequence must be carried as part of the *Hϕ* sequence or synthesized during integration.

### *HEPHAESTUS* shows evidence of horizontal gene transfer

Examination of other Eurotiales genomes in GenBank via BLAST identified a remarkably similar gene cluster in *Penicillium fuscoglaucum* strain FM041 ^33^. The shared genes between this cluster and *HEPHAESTUS* show a high level of conservation (Figure 2C) suggesting the occurrence of a horizontal gene transfer event in the evolutionary history of this region. While the left border of the *P. fuscoglaucum* cluster lies beyond the edge of the sequencing contig, the right hand terminal inverted repeat is preserved and the “ACAACTAG” motif is replaced by the similar sequence ACAAC**AT**G. Adjacent to this ACAAC**AT**G sequence are two thymine nucleotides presumably derived from an **TT**AC target site. This provides evidence that the *P. fuscoglaucum* cluster may be or have been similarly mobile.

**Fig. 2.**
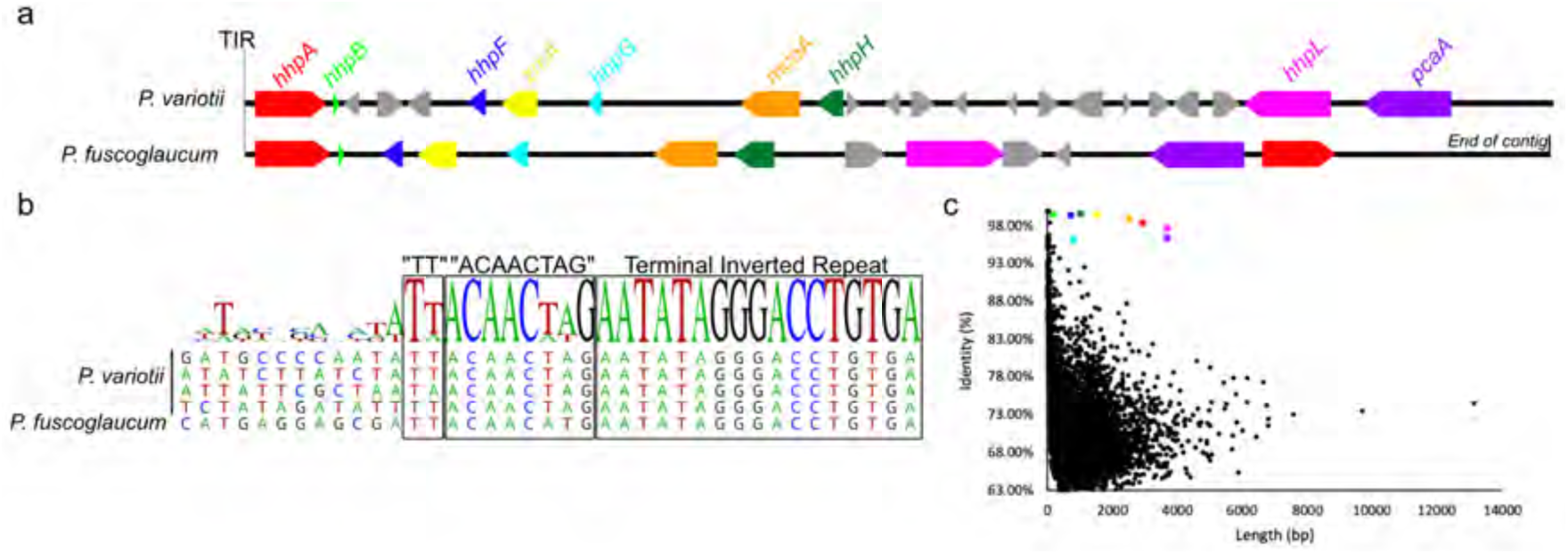
**a**, Comparison of the *Penicillium fuscoglaucum* gene cluster and *P. variotii Hϕ.* Homologous genes share the same color. **b**, The right terminal repeat is conserved in *P. fuscoglaucum* along with sequence similar to the “ACAACTAG” motif and the TT sequence presumably resulting from integration into a TTAC site. **c**, Graph of the top results of a BLAST search for all predicted *P. variotii* genes against the *P. fuscoglaucum* assembly scaffolds.Common genes between the *P. fuscoglaucum* putative metal resistance cluster and *Hϕ* (colored-coded as in panel **a**) show greater than expected conservation.

### The presence of *HEPHAESTUS* increases resistance of *P. variotii* to cadmium, lead, zinc, arsenic and copper

To test the hypothesis that the presence of *HEPHAESTUS* impacts the fitness in the presence of toxic metals we tested six *P. variotii* strains CBS 144490, FRR 1658, FRR 3823, and FRR 5287 (*Hϕ+*), and FRR 2889 and CBS 101075 (*Hϕ-*) for their ability to grow on different metal ions (Supplemental Figure 1). The *Hϕ+* strains have increased resistance to zinc, cadmium, lead and copper ions. To confirm that these resistances were conferred by *HEPHAESTUS* we examined the segregation of the resistance phenotype in the progeny of a sexual cross between two strains, CBS 144490 (*Hϕ+*) and CBS 101075 (*Hϕ-*) (Figure 3). Resistance to all four metal ions segregated with the presence of *HEPHAESTUS*. Furthermore, while characterizing resistance to arsenate we noticed a difference in early spore germination and hyphal growth between strains CBS 101075 (*Hϕ-*) and CBS 144490 (*Hϕ+*) in the presence of arsenate after 24 hours. This difference was less pronounced by four days (Figure 3).

**Fig. 3.**
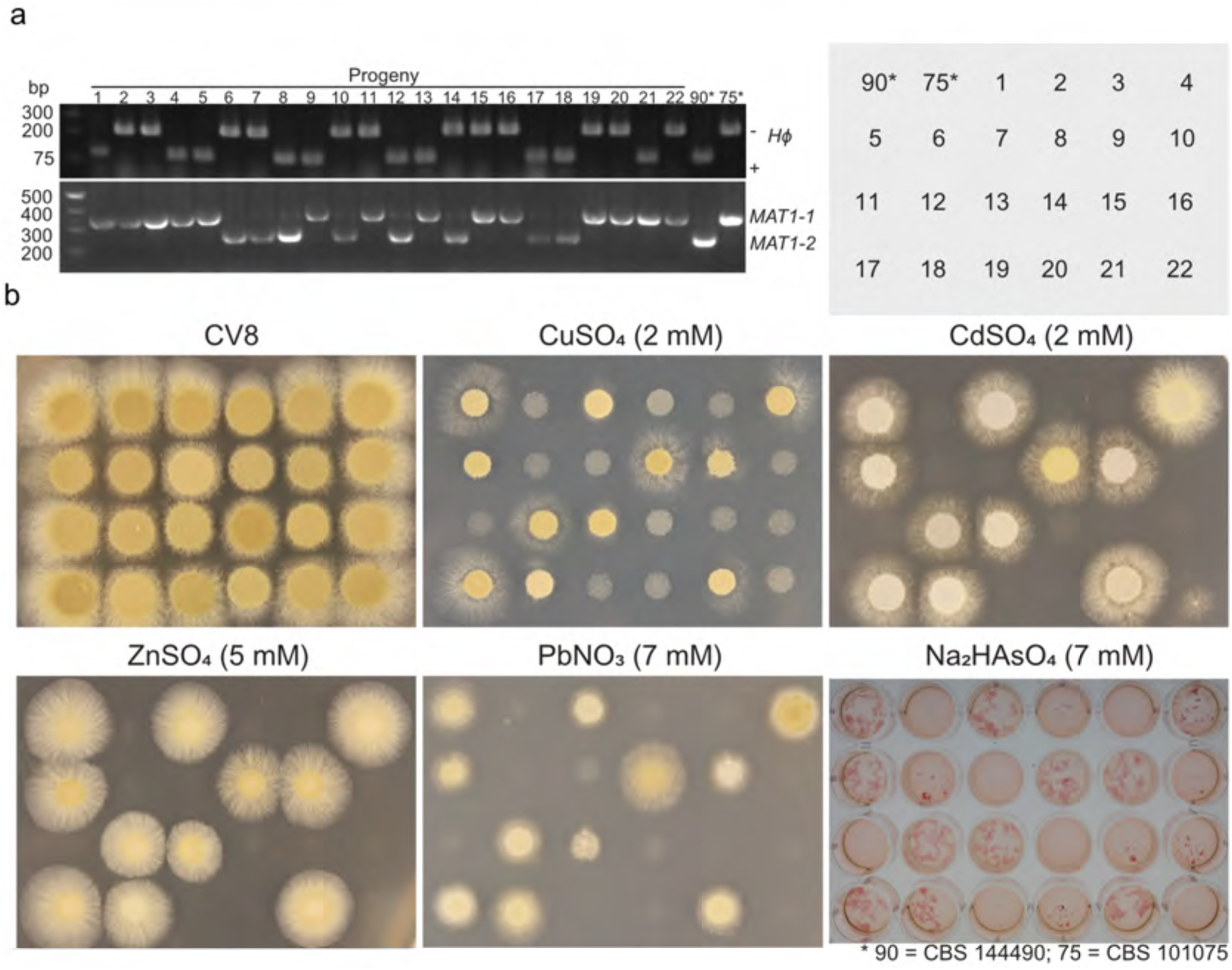
Resistance to five metal ions, Cu^2+^, Cd^2+^, Zn^2+^, Pb^2+^ and arsenate, segregates with the presence of *Hϕ* in progeny of a cross between CBS 144490 (*Hϕ+*) and CBS 101075 (*Hϕ-*). **a**, PCR analysis of the presence or absence of *Hϕ* in progeny, with the mating type locus used as a control for independent segregation. **b**, Growth at 30 °C of the two parental and 22 ascospore-derived progeny strains in cleared Campbells V8 juice plates (CV8) in presence of CuSO_4_, CdSO_4_ or ZnSO_4_ for 2 days; PbNO_3_ for 3 days. Strains were grown overnight with Na_2_HAsO_4_ in liquid media and then stained with Congo Red to increase contrast.

**Fig. S1.**
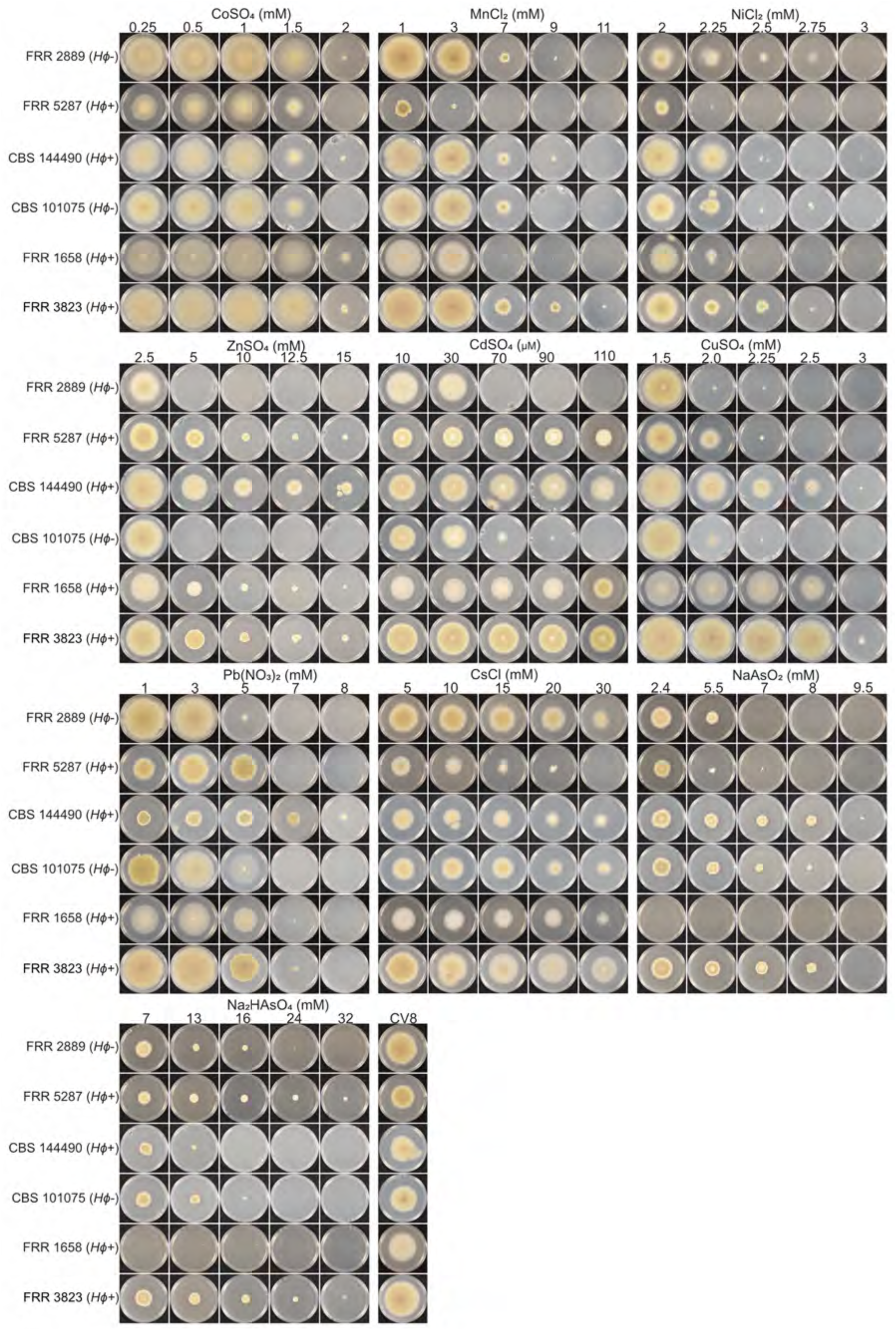
Growth for four days at 30 °C of six *P. variotii* strains on a range of concentrations of various metals. Strains with the *Hϕ* cluster are more resistant to cadmium, lead, copper and zinc than those strains lacking it.

### A P-type ATPase encoded by *pcaA* within *HEPHAESTUS* confers resistance to cadmium and lead

Having established that *Hϕ* provides resistance to toxic metals we sought to identify the specific genes conferring these resistances. The *pcaA* gene in *Hϕ* was investigated as a candidate for cadmium resistance because it shows striking sequence similarity to a previously characterized cadmium transporters in *Penicillium janthinellum* ^34^ and *Aspergillus fumigatus* ^35^. Consistent with this hypothesis, transformation of the *pcaA* gene from *Hϕ* into the *Hϕ-*strain CBS 101075 resulted in a strain with increased resistance to Cd^2+^ and Pb^2+^ (Figure 4a). Replacement of the *pcaA* coding sequence with Green Fluorescent Protein (GFP) in CBS 144490 resulted in sensitivity to cadmium. These experiments confirmed that *pcaA* in the *Hϕ* cluster is responsible for cadmium resistance.

**Fig. 4.**
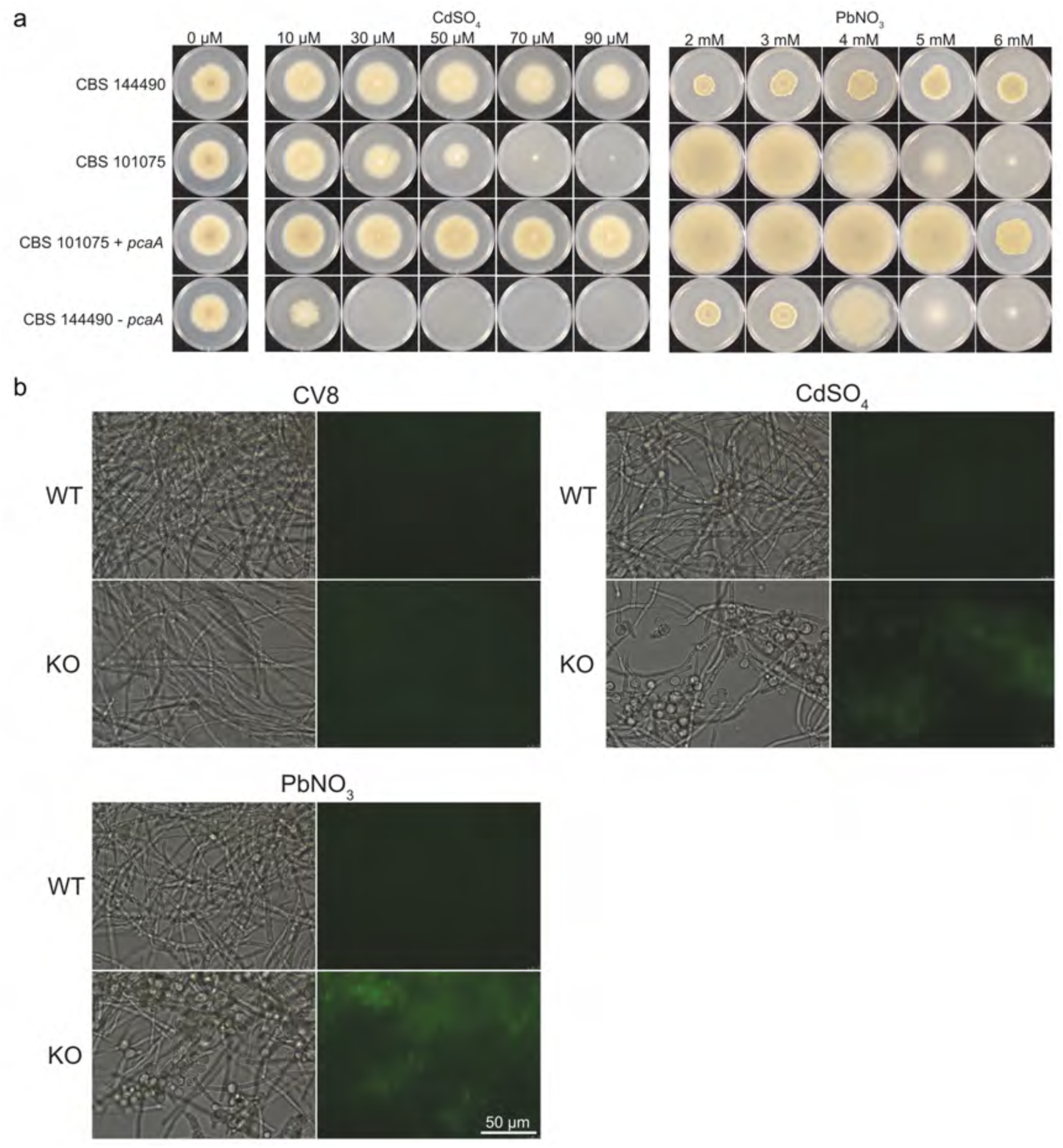
**a**, Deletion of *pcaA* in CBS14490 decreases resistance to cadmium and lead ions, and conversely transformation of *pcaA* into CBS101075 increases resistance to cadmium and lead ions demonstrating the role of *pcaA* in zinc resistance. **b**, GFP expression in the *pcaA*-GFP replacement strain increases in response to both cadmium and lead demonstrating that expression of the *pcaA* gene increases in response to high concentrations of cadmium or lead.

An unexpected finding was that in addition to conferring cadmium resistance, PcaA also confers resistance to Pb^2+^ (Figure 4a). The growth pattern differs from that of sensitivity to other metals, with rapid radial growth in strains without *Hϕ* at some concentrations of lead ions, with its role in resistance most clearly illustrated at 5 mM or higher. Previous analysis of close homologs ^34,35^ did not investigate the transport of lead; however, P-type ATPases which transport both cadmium and lead have been identified, for example AtHMA3 in the plant *Arabidopsis thaliana* ^36^. We cannot yet conclude whether the selective advantage of PcaA is primarily in response to cadmium or lead (or both).

To determine if the expression of the *pcaA* promoter in *P. variotii* CBS 144490 is regulated by these metals, a strain in which the reading frame of *pcaA* had been replaced by GFP was examined. This strain displayed green fluorescence when exposed to cadmium or lead ions, whilst low levels of fluorescence, similar to the background fluorescence of an untransformed strain, were observed in cultures without cadmium (Figure 4b).

### Zinc resistance is conferred by the *zrcA gene*

In contrast to cadmium, which is a highly-toxic non-essential metal, zinc is essential for many cellular functions ^37^. However, high levels of zinc are toxic both to humans and microorganisms ^38^. BLAST searches identified a gene, *zrcA*, with homology (36% amino acid identity) to the transporter Zrc1 that confers resistance to zinc in *Saccharomyces cerevisiae* ^39^. We investigated whether this gene was primarily responsible for the zinc resistance conferred by *Hϕ*. Transformation of *P. variotii* strain CBS 101075 with *zrcA* resulted in a strain with increased levels of resistance to zinc ions and replacement of the *zrcA* reading frame with GFP in CBS 144490 resulted in sensitivity to zinc (Figure 5a). Furthermore, the *zrcA* cDNA was able to complement the sensitivity of a *zrc1Δ cot1Δ S. cerevisiae* mutant to 5 mM ZnSO _4_ (Figure 5b). Fluorescence in the GFP replacement strain increased when zinc was added to the media, demonstrating that *zrcA* is upregulated in response to high concentrations of zinc (Figure 5c).

**Fig. 5.**
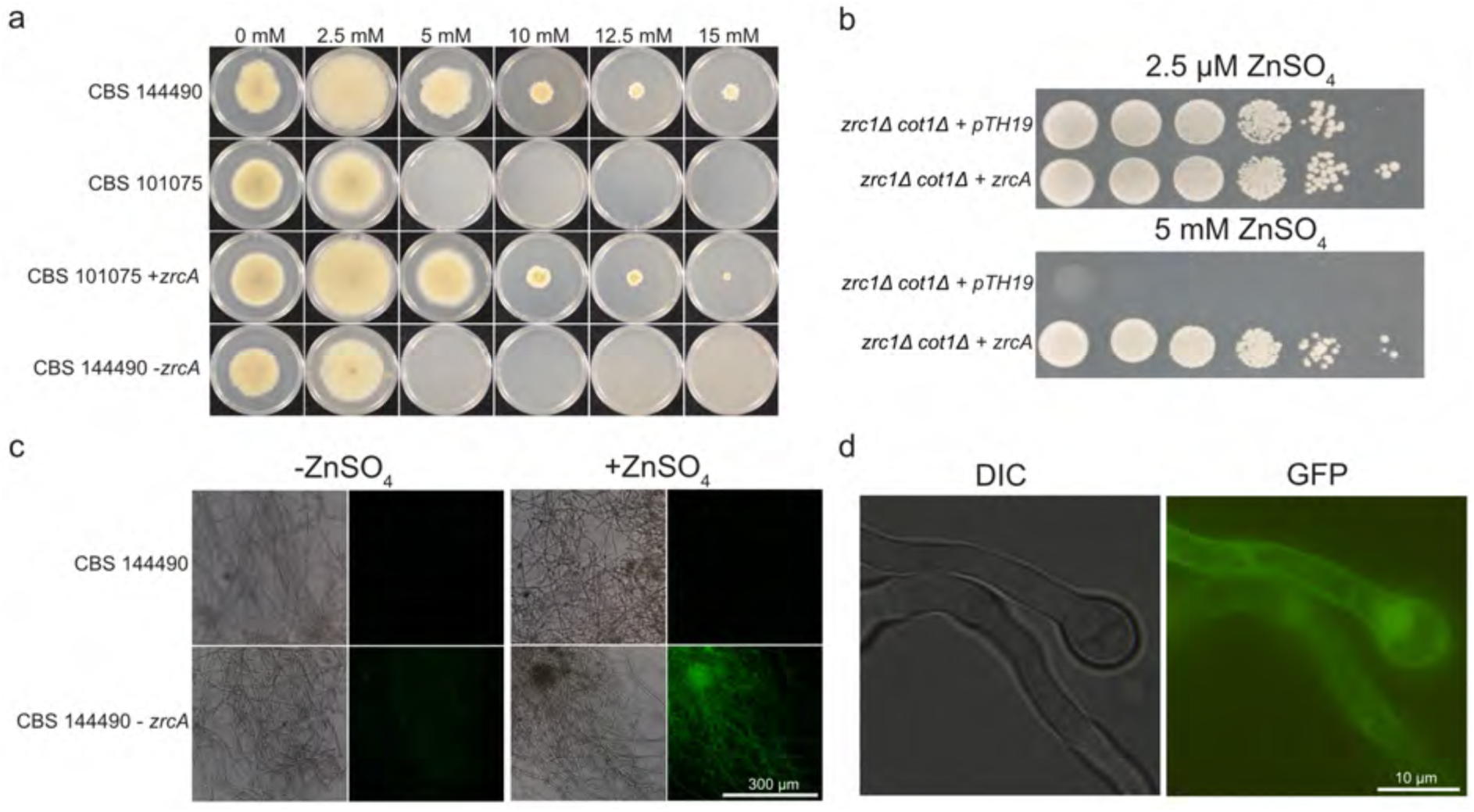
**a**, Deletion of *zrcA* in CBS 144490 decreases resistance to zinc ions and addition of *zrcA* to CBS 101075 increases resistance to zinc ions. **b**, A *zrcA*-containing construct complements the zinc sensitivity of a *zrc1Δ cot1Δ S. cerevisiae* mutant, unlike the empty vector (pTH19). **c**, GFP expression in the *zrcA*-GFP replacement strain increases in response to zinc demonstrating that expression of the *zrcA* gene is increased in response to high concentrations of zinc ions. **d**, A *zrcA*-GFP fusion protein localizes largely to the cell membrane indicating a possible role in export of zinc from the cell.

A strain of *P. variotii* in which GFP was fused behind ZrcA and expressed under the constitutive *Leptosphaeria maculans* actin promoter was developed to investigate the subcellular localization of ZrcA. This strain showed GFP fluorescence consistent with plasma membrane localization, as well as within the vacuole (Figure 5d). This contrasts to the known localization of Zrc1 homologs. In *S. cerevisiae* and *Cryptococcus neoformans* Zrc1 is localized to the vacuolar membrane and leads to the accumulation of zinc in the vacuole ^40,41^. However, in *Candida albicans* and *Hebeloma cylindrosporum* the Zrc1 homologs appear to be localized to the endoplasmic reticulum and leads to zinc storage in so called “Zincosomes” ^42,43^. The presence of *P. variotii* ZrcA in this membrane suggests a possible role in exporting zinc ions out of the cell rather than in involvement in intracellular zinc storage as in the case of Zrc1 in *S. cerevisiae, C. albicans* and *H. cylindrosporum*.

### A multigene region of *Hϕ* confers arsenic resistance

A sub-region of *Hϕ* contains a cluster of genes putatively involved in arsenic resistance, including homologs of *arsH* ^44^, *arsC* ^45^, and *arsB* ^46^ as well as genes encoding a cyclin similar to Pho80 ^47^ and an arsenic methyltransferase ^48^ (Figure 6a). Deletion of a 9,478 bp region containing most of these genes resulted in sensitivity to arsenic and complementation of this mutant with these genes restored arsenic resistance (Figure 6b). The key gene(s) responsible for the increase in arsenic tolerance associated with *Hϕ* within this region have yet to be identified and it is possible that some of these genes act redundantly or synergistically. In addition to the genes in the cluster, other copies of a subset of these genes are found in the *P. variotii* genome. Further studies of this region in *P. variotii* or other species will provide greater insight into the mechanisms of arsenic resistance in filamentous fungi.

**Fig. 6.**
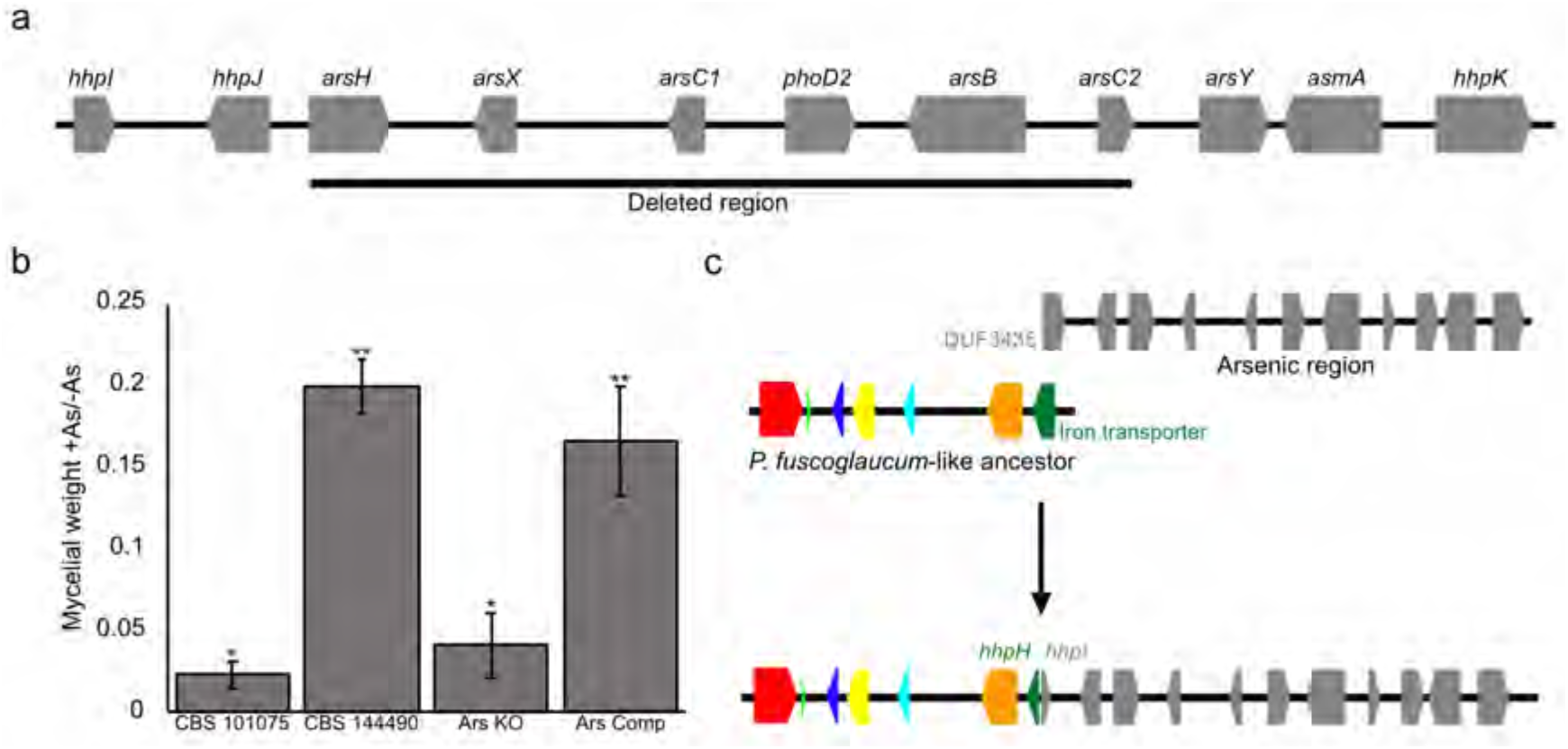
**a**, The putative arsenic-associated region contains 10 genes (in addition to *hhpI*, which is possibly a pseudogene) in *P. variotii*. A sub-region containing six of these genes was deleted by homologous recombination. **b**, This deletion resulted in strains (Ars KO) with significantly reduced growth (mycelial weight) in 7 mM arsenate after 48 h. Complementation of this mutant (Ars Comp) restored arsenic resistance. **c**, Model explaining the origin of the arsenic region in *Hϕ* as an insertion of a preformed arsenic cluster into a *Penicillium fuscoglaucum* cluster-like ancestor.

One explanation for the presence of the genes for arsenic detoxification within *Hϕ* is insertion of the region into an ancestor sequence similar to the *Penicillium fuscoglaucum* cluster, which lacks these genes. This hypothesis is supported by the presence of two putative pseudogenes that have been created at the right junction of this hypothesized insertion. The first is *hhpH*, which is a truncated putative iron transporter. The *P. fuscoglaucum* cluster contains a full-length homolog spanning across this junction, suggesting that the arsenic region was mostly likely inserted into the ancestral cluster rather than deleted from it. The second pseudogene is *hhpI*, which is a truncated homolog of a conserved group of proteins containing a domain of unknown function (DUF3435). A model describing how such a junction may have formed is presented in Figure 6c.

### The multicopper oxidase McoA confers resistance to copper

Despite the important use of copper-based fungicides for over 100 years, knowledge of copper resistance in filamentous fungi is limited. We hypothesized that the *mcoA* gene encoding a putative multicopper oxidase within the *HEPHAESTUS* region might confer resistance to copper ions, since other multicopper oxidases confer resistance to copper ions in both bacteria (e.g. CueO in *Escherichia coli* ^49^) and fungi (e.g. Fet3 in *S. cerevisiae* ^50^). Consistent with this hypothesis, transformation of CBS 101075 with *mcoA* conferred resistance to copper ions (Fig. 7a). Furthermore, qPCR analysis showed that expression of *mcoA* in 1.5 mM copper was 4-5 times higher than when strain CBS 144490 was grown without copper (Fig. 7b). This is the first multicopper oxidase conferring resistance to copper reported from a filamentous fungus.

**Fig. 7.**
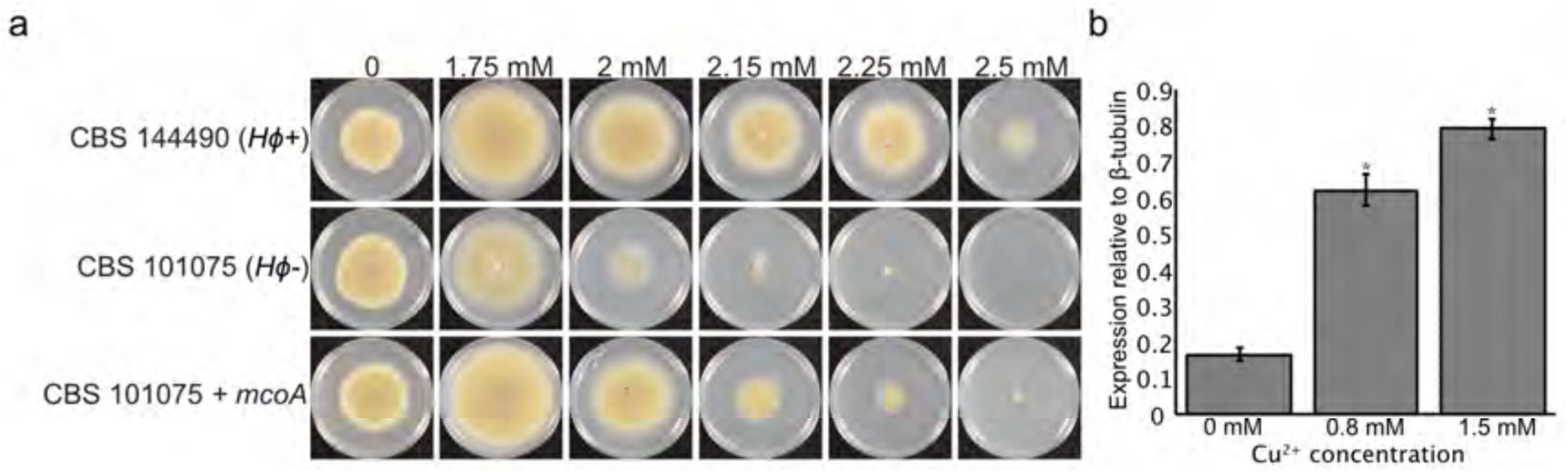
The multicopper oxidase McoA encoded within *Hϕ* confers resistance to copper ions. **a**, Transformation of *mcoA* into CBS 101075 (*Hϕ-*) provides increased resistance to copper, suggesting that McoA contributes to the copper resistance conferred by *Hϕ*. **b**, qPCR analysis shows increased expression of *mcoA* in the presence of copper in strain CBS 144490. *Differences between expression at different copper concentrations are statistically significant (P<0.01) as assessed using a two-tailed t-tests assuming unequal variance. Independent biological replicates were measured, 0 mM and 1.5 mM n=4; 0.8 mM n=3. Error bars represent plus or minus one standard deviation from the mean.

## DISCUSSION

Most cases of horizontal DNA transfer in eukaryotes involve transposons ^51^. Transposons have been widely characterized in eukaryotes since their discovery in maize 70 years ago ^52^. However, the potential benefits of genes carried within them to the host are not usually apparent. One exception is a putative transposon, ToxhAT, involved in the movement of the ToxA host specificity toxin between plant pathogens ^53^. ToxhAT contains 14 kb of DNA flanked by TIRs and appears to have horizontally transferred as a transposon between *Parastagonospora/Pyrenophora* and *Bipolaris*. However recent mobility in the *Bipolaris sorokiniana* genome appears to be as part of a broader genomic island rather than through its own transposition. An important difference between ToxhAT and *Hϕ* is that a single gene within ToxhAT, namely ToxA, confers the phenotype as opposed to a multitude of genes. Recently, increasingly large transposable elements have been found in eukaryotes. These include helitrons in maize capable of capturing up to nine gene fragments and totaling up to 39 kb ^54^, and the tetratorn elements of fish that are well over 150 kb ^21^. The discovery of these large transposable elements in eukaryotes has prompted the hypothesis that such transposons might be capable of carrying greater amounts of useful “cargo” including entire gene clusters ^22^.

Here we describe *Hϕ*, a large (∼85 kb) region in a fungal genome that contains a number of genes involved in resistance to toxic metal ions. These genes include a P-type ATPase conferring resistance to cadmium and lead, a *zrc1* homolog conferring resistance to zinc, a multicopper oxidase conferring resistance to copper and subregion of the cluster conferring resistance to arsenic compounds. Further, we present evidence that *Hϕ* is a transposable element. The four *Hϕ*+ strains sequenced each contain *Hϕ* at a unique genomic location on separate assembly contigs demonstrating mobility within the *P. variotii* genome (Figure 1c). Moreover, examination of the flanking sequences of *Hϕ* suggests an active mechanism of transposition given the perfect preservation of the *Hϕ* border sequences (including terminal inverted repeats) and preference for particular target sites (Figure 1d). Often, transposition of DNA transposons involves target site duplication ^55^. *Hϕ* does not appear to employ such duplications but rather makes use of microhomology to the target site. This is reminiscent of Crypton transposons, which also make use of microhomology to the target site ^56^. However, Cryptons differ from *Hϕ* in key respects including that they lack TIRs. There is also precedent for an absence of target site duplication, namely the *Spy* transposons ^57^. Another distinctive feature is that *Hϕ* carries an “ACAACTAG” motif outside one of the terminal inverted repeats; we are not aware of a similar feature in other transposons.

*Hϕ* carries genes with homology to those involved in transposition, including a putative Tc-1 like transposase, and a reverse transcriptase homolog, which may play a role in the movement of *Hϕ*, as might the pair of highly conserved internal repeats which initially frustrated DNA sequencing assembly. While the mechanisms of *Hϕ* transposition will require further studies to illuminate them, *Hϕ* is likely capable of transposition by a specific and possibly novel mechanism given that the integration sites do show a consistent pattern different from known transposons. At 85 kb *Hϕ* represents an exceptionally large fungal transposon (although smaller than the Teratorn element in fish ^21^) and carries substantially more host beneficial genes than any previously described transposable element. Other examples of beneficial gene clusters in fungi (including those involved in secondary metabolite synthesis) have also been hypothesized to move via horizontal gene transfer ^16^ but do not show evidence of transposition within the genome.

A comparable region encoding for metal resistance has not previously been described in fungi, with *Hϕ* being set apart both in the number of genes encoded (> 30), the number of metal ion resistances conferred (≥ 5), and evidence of mobility. However, there are parallels with bacterial operons for metal ion detoxification. For instance, ICEs can contain large sets of genes that are involved in the detoxification of multiple metal ions, and such elements are able to spread through replication as plasmids and via conjugation with other bacteria ^9,10,58^.

Evidence for a past horizontal DNA transfer event in the evolutionary history of the *Hϕ* cluster is provided by the strikingly high nucleotide similarity between elements of this cluster and a similar, albeit incomplete, cluster in *P. fuscoglaucum.* Curiously, *P. fuscoglaucum* is from a group of fungi noted for a high level of HGT ^16,59,60^. The high level of nucleotide identity between the two clusters (∼95%) is inconsistent with vertical gene transmission from an ancient common ancestor. We thus suggest that *HEPHAESTUS* most likely represents a recent horizontal DNA transfer event conferring resistance to various metals in certain *P. variotii* strains. This implies that as in the case of prokaryote ICEs ^9-11^, HGT is likely to play a role in evolutionary adaptation to metal stress in fungi.

The *P. variotii* strains in this study were not accompanied by detailed collection data so we were unable to draw correlations between the presence of *HEPHAESTUS* and the ecological source of the strains. However, given the seemingly diverse and poorly understood ecological niches which *P. variotii* appears to fill, an important subject for future studies will be the distribution of the cluster from a population biology perspective in a greater number of strains.

An implication of carrying host beneficial genes on a mobile element in *P. variotii* is that we expect that such regions will be subjected to Repeat Induced Point (RIP) mutation should they become duplicated in the *P. variotii* genome ^29^. RIP is a fungal defense against transposable elements that inactivates duplicated DNA during sexual reproduction through C to T mutations ^61^. Thus, we predict that the “domestication” of these genes in a stable genomic location may be favored. This selective pressure may limit the life cycle of positively selected transposons. Supporting this is the presence of a P-type ATPase in *Aspergillus clavatus* strain NRRL 1 with unusually high homology (93% nucleotide identity) to the *pcaA* gene of *Hϕ* (GenBank XM_001276247.1 ^62^) yet not located in a similar genomic context. We speculate that this gene may represent a domesticated relic of an element similar to *Hϕ*. If these elements are not evolutionarily stable, then the window of time in which they can be observed may be narrow and thus their importance as drivers of evolutionary change obscured.

In summary, we have identified a new mobile element in fungi involved in tolerance to toxic metal ions. This region contains over 30 genes many of which appear to have potential roles in metal detoxification, some of which we have characterized in this study. *Hϕ* is the first eukaryotic transposable element found to carry such a significant complement of host-beneficial genes and highlights the possible role of large transposable elements in the horizontal transfer of large genomic regions between eukaryotes. Future studies should explore its origins in related Eurotiales species and the mechanism by which such a large region is able to transpose.

## METHODS

### DNA sequence refinement and annotation of the *Hϕ* region

Examination of the next generation sequencing reads of *P. variotii* CBS 144490 assembled to contig 123, where the *Hϕ* gene cluster is located, revealed a number of regions of ambiguity. Furthermore, comparison of the genes by BLAST against fungal genome sequences in GenBank identified a related cluster found in the cheese-associated species *Penicillium fuscoglaucum* ^33^, whose arrangement of genes also included homologs in *P. variotii* found on contig 14, which is another region found in strain CBS 144490 and not CBS 101075. Hence, specific regions were amplified by PCR and sequenced to provide a high confidence level of assembly, which joins contig 14 into the middle of the cluster on contig 123. The reason that this was not assembled by the program Velvet is likely due to ∼300 bp of perfectly repeated DNA flanking this region. The compete *Hϕ* sequence from strain CBS 144490 has been deposited to GenBank (accession MT022027). Wild type strains can be obtained from culture collections and genetically manipulated strains can be provided by the authors on request.

### Strains and culturing

The two *P. variotii* strains with their genomes sequenced are CBS 101075 and CBS 144490; these strains were routinely cultured on cleared Campbell’s V8 juice (CV8) agar. Four additional strains FRR 1658 (*Hϕ*+, ITS GenBank MN880081), FRR 2889 (*Hϕ*-, ITS GenBank MN880082), FRR 3823 (*Hϕ*+, ITS GenBank MN880083) and FRR 5287 (*Hϕ*+, ITS GenBank MN880084) were obtained from the FRR culture collection maintained by CSIRO (North Ryde, New South Wales, Australia). Correct identification of the four FRR strains was confirmed using an ITS phylogenic tree (Supplementary Figure 2) generated in MrBayes ^63^ based on a previous taxonomic study of the *Paecilomyces* genus ^64^.

*Escherichia coli* strain NEB® 5-alpha was used for propagation of plasmid DNA. *Agrobacterium tumefaciens* strain EHA105 was used for transformation of *P. variotii*. The bacterial strains were maintained on Luria broth (LB) agar, supplemented with either kanamycin (50 mg/L) or ampicillin (50 mg/L) in plasmid-carrying strains. *Saccharomyces cerevisiae* strain BY4742 and derivatives was used for *zrcA* complementation experiments. *S. cerevisiae* was cultured on yeast extract peptone dextrose (YPD), YPD +G418 (for selection of the KanMX construct), yeast nitrogen base (YNB) -leucine (for selection of the *leu2* construct) or YNB -uracil (for selection of the pTH19 plasmid).

**Fig. S2.**
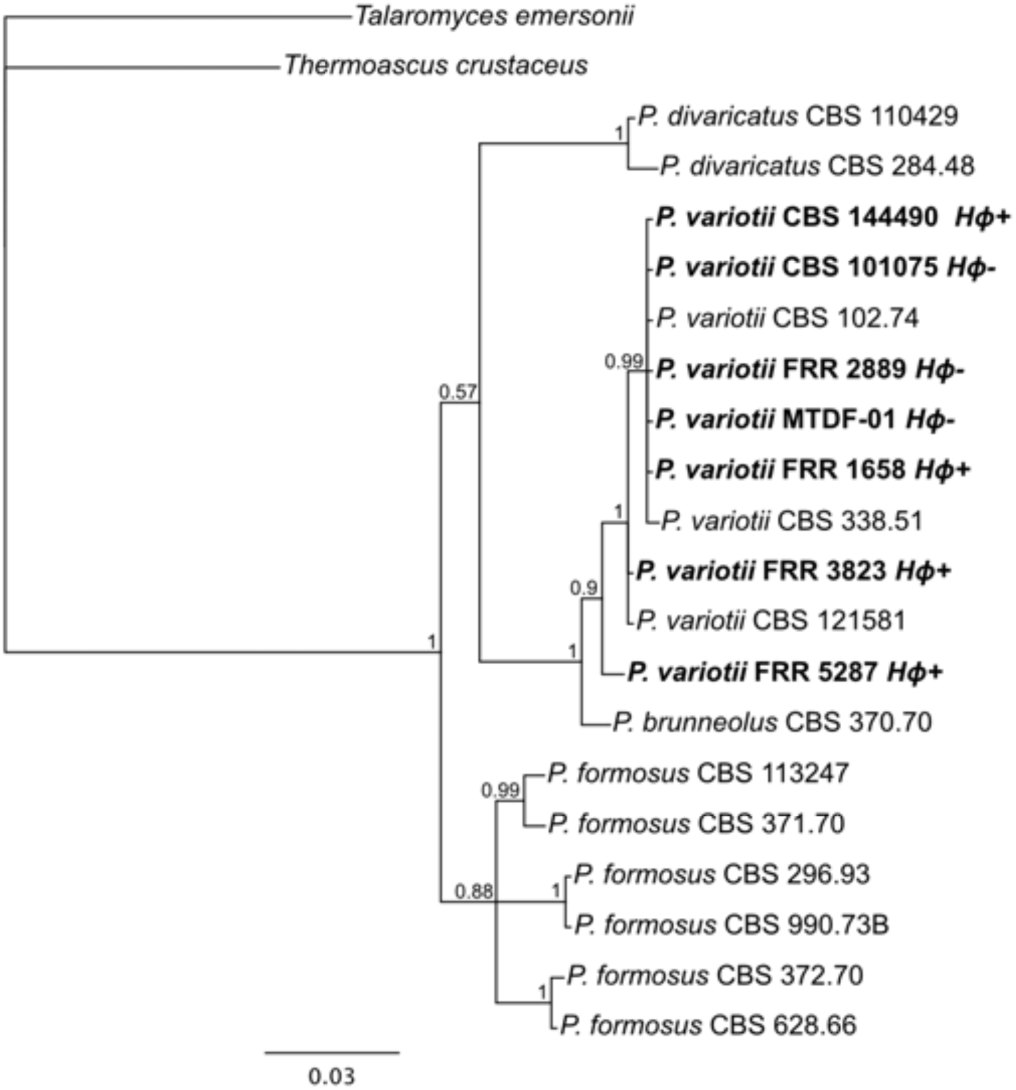
Phylogeny of *Paecilomyces* strains based on the ITS region, illustrating that four of the FRR culture collection strains are *P. variotii*. Strains in which the presence or absence of *Hϕ* is known are in bold font. Posterior probabilities are given for each clade.

### Genetic segregation

Crosses between strains CBS 101075 and CBS 144490 were conducted as previously described, and 22 ascospore-derived colonies obtained ^24,29^. The mating type of each progeny was determined by PCR using primers MAI0448, MAI0449, MAI0450 and MAI0451. The presence or absence of *Hϕ* was assessed using primer pairs AP89 and AP90, specific for *Hϕ*, and AP92 and AP93, which amplify across the *Hϕ* insertion site. Primer sequences are provided in Table S2.

### Cloning and DNA manipulations

The *pcaA* and *zrcA* open reading frames were amplified with primer pairs AP104 and AP105 and AP297 and AP298, and cloned into plasmid PLAU2 linearized with BglII, which enables constitutive expression from the actin promoter of *Leptosphaeria maculans* ^29,65^. A construct expressing a ZrcA-GFP fusion protein was generated by amplifying the *zrcA* reading frame with primers AP351 and AP352, and the *GFP* reading frame with primers AP353 and AP354 and cloning into the BglII site of plasmid PLAU2.

Constructs for replacement of the *pcaA* and *zrcA* genes with Green Fluorescent Protein (GFP) via homologous recombination were made by PCR amplification of the required gene regions followed by cloning of these PCR products into plasmid pPZP-201BK ^66^ linearized with EcoRI and HindIII. For *pcaA* the regions of homology were amplified using primer pairs AP219 and AP220 and AP223 and AP224. The *GFP*-*hph* region was amplified from a GFP-tagging construct generated previously ^67^ using primers AP221 and AP222. For *zrcA* the regions of homology were amplified using primer pairs AP299 and AP302 and AP305 and AP307. The *GFP-hph* was amplified using primers AP303 and AP304.

A construct for the deletion of a 9,478 bp region containing genes putatively involved in arsenic resistance was similarly constructed using the NEBuilder assembly kit (New England Biolabs, Ispwich, MA, USA). Genomic DNA on the 5’ flank of the region was amplified using primers AP34 and AP35, and on the 3’ flank with primers AP36 and AP37. The *hph* expression cassette conferring hygromycin resistance was amplified with primers AP63 and AP64. The three fragments were assembled into plasmid pPZP-201BK linearized with EcoRI and HindIII. A construct to complement this deletion was generated by amplifying the required region in 4 pieces by PCR using primer pairs AP308 and AP309, AP310 and AP314, AP315 and AP311 and AP312 and AP313. These fragments were cloned into plasmid pMAI2 ^65^ linearized with XbaI to recreate a 12,176 bp fragment of the genome.

### Transformation of *P. variotii* and screening for correct integration events

Donor DNA was cloned into T-DNAs and transformed via *Agrobacterium tumefaciens* into the genome of *P. variotii*, as described previously ^29^.

In cases where targeted integration of the construct was required, DNA was prepared using a CTAB buffer protocol ^68^ and correct integration of the construct was assessed by PCR. For *pcaA*, primer pairs AP235 and AP251 and AP236 and AP252 were used. For *zrcA*, primer pairs AP323 and AP251 and AP324 and AP252 were used. Deletion of the arsenic region was assessed using primers pairs AP116 and MAI0023 and AP117 and MAI0022.

### Yeast complementation

Homologous recombination using a PCR product was used to disrupt the *ZRC1* and its paralog *COT1* in *S. cerevisiae*. In the case of *ZRC1*, the *LEU2* allele was amplified using AP316 and AP317 off the plasmid pGAD ^69^ with appropriate homologous sequence added the primers for targeting to the *ZRC1* locus. In the case of *COT1*, primers AP327 and AP328 were used to amplify the KanMX expression cassette from plasmid pFA6a-GFP(S65T)-kanMX6 ^70^ with appropriate homologous sequence added the primers for targeting to the *COT1* locus. These PCR products were sequentially transformed as described previously ^71^ into *S. cerevisiae* strain BY4742, selecting on either media lacking leucine in the case of *LEU2* or with 200 µg/ml G418 in the case of KanMX. DNA was prepared from the transformants ^72^ and correct integration of the deletion construct was assessed by PCR using primers AP325 and AP326 for *ZRC1*, and AP341 and AP342 for *COT1*.

The *zrcA* cDNA was cloned into plasmid pTH19 to express ZrcA under the control of the constitutive alcohol dehydrogenase 1 promoter in *S. cerevisiae* ^73^, as follows. RNA was extracted using Trizol and treated with DnaseI (New England Biolabs). The RNA was reverse transcribed using AMV reverse transcriptase (New England Biolabs) and an oligo-dT primer. The Zrc1 cDNA was amplified using Q5 Polymerase Master Mix (New England Biolabs) with primers AP337 and AP340. The PCR product was then cloned into pTH19 linearized with BamHI and HindIII using the NEBbuilder assembly kit (New England Biolabs). The plasmid was transformed into yeast as described previously ^71^ selecting for transformants on media lacking uracil.

### Quantitative PCR

RNA was extracted using Trizol from 48 hours old cultures grown in 10% Campbells V8 juice supplemented with different concentrations of copper sulfate. The RNA was reverse transcribed using AMV reverse transcriptase (New England Biolabs) using oligo-dT. The *mcoA* gene was amplified using primers AP369 and AP370 and the constitutive beta-tubulin gene was amplified as described previously ^67^.

### Microscopy

Strains were examined under a Leica DM6000 B microscope equipped with a Leica DFC 450 C camera. GFP fluorescence was visualized using the “I3” filter cube.

### Genome sequencing and assembly

Genomic DNA was extracted from stationary liquid cultures of *P. variotii* strains FRR 1658, FRR 2889, FRR 3823 and FRR 5287, as described previously ^68^. Genomes were sequenced using an Illumina NovaSeq 6000 System at the Murdoch Children’s Research Institute (Melbourne, Australia). Reads were trimmed using BBduk and mapped to the *Hϕ* region ^74^ using Geneious version 11.1.5 to identify junctions. Raw reads have been deposited in the GenBank SRA database (BioProject ID: PRJNA604095).

## ACKNOWLEDGEMENTS

We thank the University of Melbourne and Australasian Mycological Society for support for this research, Mark Wilson (CSIRO) for providing fungal strains, and Barbara Howlett for comments on the manuscript.

---

**Table S1.** Genes present within the *Hϕ* region.

| Protein ID in JGI | Gene name | Putative function |
| --- | --- | --- |
| 195182 | *hhpA* | DUF3435 |
| 167128 | *hhpB* | Transcription factor, MADS-box. Possible pseudogene. |
| 290670 | *hhpC* | Reverse transcriptase |
| 276811 | *hhpE* | Mitochondrial Fe<sup>2+</sup> transporter MMT1 |
| NA |  | Tn pseudogene |
| 232099 | *hhpF* |  |
| 179937 | *zrcA* | Zinc transporter |
| 232097 | *hhpG* |  |
| 276812 | *mcoA* | Multicopper oxidase |
| 290673 | *hhpH* | Low affinity Fe<sup>2+</sup> transporter |
| 290674 | *hhpI* | Truncated DUF3435 domain protein. Possible pseudogene. |
| 290675 | *hhpJ* |  |
| 276813 | *arsH* | NADPH-dependent FMN reductase-like<br>arsH |
| 239850 | *arsX* | Unknown gene conserved in ars clusters |
| 239851 | *arsC1* | Arsenate reductase, arsC |
| 239852 | *phoD2* | Cyclin, PHO80 |
| 290679 | *arsB* | Arsenite efflux transporter |
| 239854 | *arsC2* | Arsenate reductase |
| 290681 | *arsY* | Intermediate filament Protein. Next to Cyt19 in other fungal genomes. |
| 290682 | *asmA* | Arsenic methyltransferase; like Cyt19 |
| 290683 | *hhpK* | K<sup>+</sup>-dependent Ca<sup>2+</sup>/Na<sup>+</sup> exchanger |
| 239858 | *hhpL* | Metal transporting P-type ATPase |
| 195275 | *pcaA* | Metal transporting P-type ATPase |
| 276819 | *hhpM* | Transcription factor, MADS-box |
| 276820 | *hhpN* | Tc1 transposon |
| 290686 | *hhpO* |  |
| 290687 | *hhpP* |  |
| 290688 | *hhpQ* |  |
| 195756 | *hhpR* | metalloprotease |
| 290691 | *hhpS* | AAA ATPase; mitochondrial chaperone BCS1 |
| 290692 | *hhpT* |  |
| 290693 | *hhpU* |  |
| 290694 | *hhpV* |  |
| 239869 | *hhpW* |  |
| 239870 | *hhpX* |  |

**Table S2.**
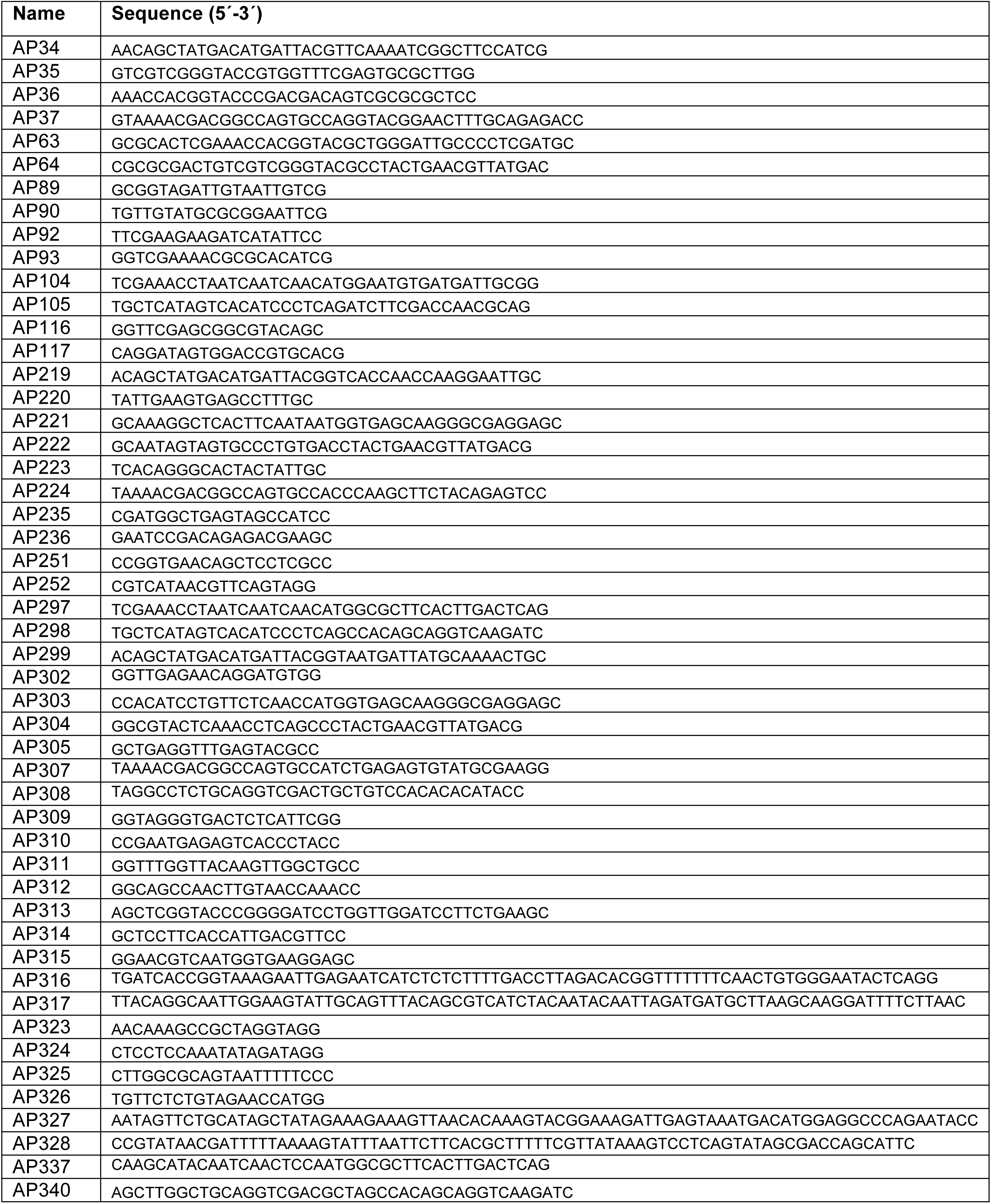
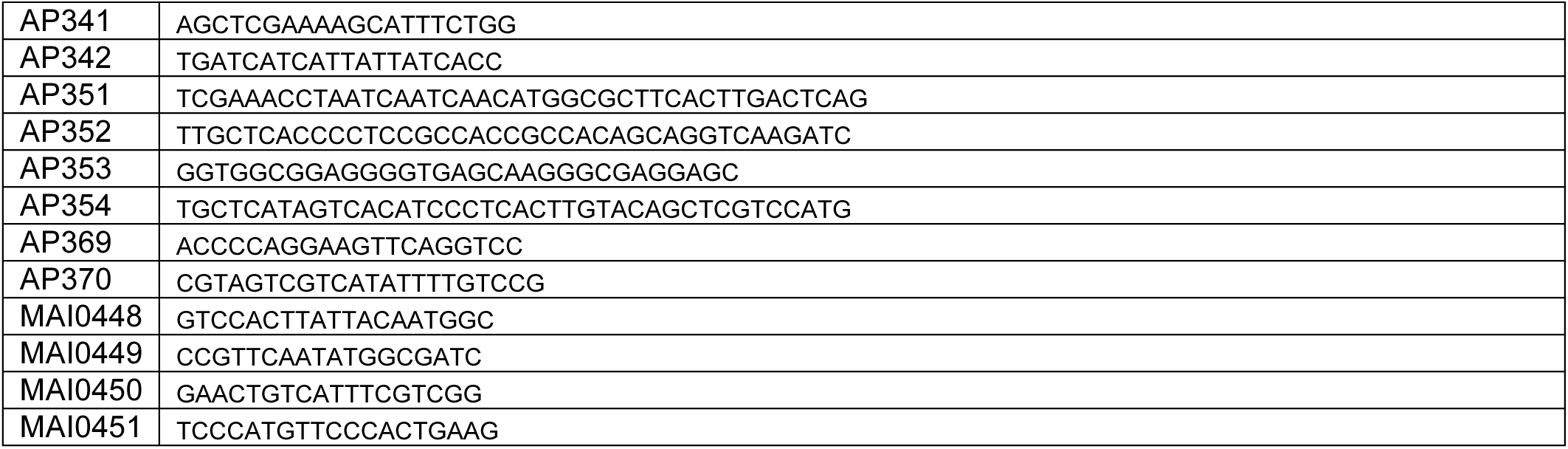
primers used in this study.

